# A Micro-Engineered Airway Lung-Chip Models Key Features of Viral-Induced Exacerbation of Asthma

**DOI:** 10.1101/2020.02.02.931055

**Authors:** J. Nawroth, C. Lucchesi, D. Cheng, A. Shukla, J. Ngyuen, T. Shroff, K. Karalis, H-H. Lee, S. Alves, G. A. Hamilton, M. Salmon, R. Villenave

## Abstract

**Rationale:** Viral-induced exacerbation of asthma remain a major cause of hospitalization and mortality. New human relevant models of the airways are urgently needed to understand how respiratory infections may trigger asthma attacks, and to advance treatment development.

**Objectives:** To develop a new human relevant model of rhinovirus-induced asthma exacerbation that recapitulates viral infection of asthmatic airway epithelium, neutrophil transepithelial migration, and enables evaluation of immunomodulatory therapy.

**Methods:** A micro-engineered model of fully differentiated human mucociliary airway epithelium was stimulated with IL-13 to induce a Th2-type asthmatic phenotype and infected with live human rhinovirus 16 (HRV16) to reproduce key features of viral-induced asthma exacerbation.

**Measurements and Main Results:** Infection with HRV16 replicated key hallmarks of the cytopathology and inflammatory responses observed in human airways. Generation of a Th2 microenvironment through exogenous IL-13 stimulation induced features of asthmatics airways, including goblet cell hyperplasia, reduction of cilia beating frequency, and endothelial activation, but did not alter rhinovirus infectivity or replication. High resolution kinetic analysis of secreted inflammatory markers revealed that IL-13 treatment altered the IL-6, IFN-λ1, and CXCL10 secretion in response to HRV16. Neutrophil transepithelial migration was greatest when viral infection was combined with IL-13 treatment, while treatment with MK-7123, a CXCR2 antagonist, reduced neutrophil diapedesis in all conditions.

**Conclusions:** This micro-engineered Airway Lung-Chip provides a novel human-relevant platform for exploring the complex mechanisms underlying viral-induced asthma exacerbation. Our data suggest that IL-13 may impair the hosts’ ability to mount an appropriate and coordinated immune response to rhinovirus infection. We also show that the Airway Lung-Chip can be used to assess the efficacy of modulators of the immune response.

**Note:** Emulate®, Human Emulation System®, S-1™, ER-1™, and ER-2™ are trademarks of Emulate, Inc., and any other trademarks used herein remain with their respective holders. The technology disclosed in this document may be covered by one or more patents or patent applications, and no license to these is granted herein. You are solely responsible for determining whether you have all intellectual property rights that are necessary for your intended use of any of the disclosed materials, and whether you are required to obtain any additional intellectual property rights from a third party. Further information is available by contacting the authors.

**At a Glance Commentary:** *Scientific Knowledge on the Subject:* New therapies for asthma exacerbations remain a significant unmet medical need. Development of human relevant preclinical models are needed to further elucidate the complex mechanisms underlying asthma exacerbation and investigate new therapeutic strategies.

*What This Study Adds to the Field:* Using a human Airway Lung-Chip model, we show here for the first time a live human rhinovirus (HRV) infection of the asthmatic epithelium that recapitulates complex features of viral-induced asthma exacerbation. The dynamic microenvironment of the chip enables the real-time study of virus infection, epithelial response, and immune cell recruitment under healthy and asthmatic conditions. The model reproduces key endpoints that have been observed in asthmatics and individuals infected with rhinovirus including the ciliated cell sloughing, altered cilia beating frequency, goblet cell hyperplasia, increased expression of adhesion molecules in microvascular endothelial cells, and inflammatory mediator release. High-resolution temporal analysis of secreted inflammatory markers enabled by dynamic sampling revealed alteration of IL-6, IFN-λ1 and CXCL10 secretory phases after rhinovirus infection in an IL-13 high environment. Leveraging high-content imaging and analysis of circulating inflammatory cells, we demonstrated the efficacy of a CXCR2 antagonist to reduce adhesion, motility, and transmigration of perfused human neutrophils. Thus, this micro-engineered chip may offer a powerful addition to preclinical models for understanding mechanisms underlying asthma exacerbation pathology and developing new therapeutic strategies.

## Introduction

Asthma is a chronic inflammatory disease of the respiratory tract defined by shortness of breath, wheezing, and chest tightness that affects 300 million people worldwide^1^. In most cases the disease can be managed with inhaled corticosteroids and bronchodilators; however, up to 1/8 of patients experience asthma exacerbations that can lead to emergency hospitalizations and death^2^. Asthma exacerbations are characterized by acute episodes of decreased lung function and result in significant morbidity, mortality and economic costs^3,4^. Prevention and treatment of asthma attacks hence remain an area of considerable unmet medical need^5^. A major trigger of asthma attacks is human rhinovirus (HRV) infection of asthmatic airways^6–8^. New human-relevant models are critically needed to study how a respiratory infection may provoke asthma attacks, and to advance the development of therapeutics^9^. To address this challenge, we leveraged a human Airway Lung-Chip featuring mucociliary airway epithelium and hemodynamic perfusion^10^ to model key elements of viral-induced exacerbation of asthma. Our platform recapitulates (1) the asthmatic remodeling of human mucociliary airway epithelium, such as goblet cell hyperplasia^11^, (2) cytopathic effects and epithelial inflammation in response to HRV infection^12^, and (3) recruitment of immune cells to the epithelium^13^. We show that in conjunction with innovative high-content imaging, microfluidic sampling, and compound testing, the Airway Lung-Chip helps to elucidate and probe the pathological mechanisms that arise from the interaction of HRV infections with asthmatic airways, including impaired interferon responses and altered immune cell transmigration. Taken together, we demonstrate that our human Airway Lung-Chip provides a powerful new platform for studying the time course, mechanisms and potential treatments of viral-induced asthma exacerbations.

## Methods

Complete methods can be found in the online supplement.

### Cell differentiation and HRV infection on-chip

Cells were cultured and differentiated as previously described ^10^. Briefly, human primary airway epithelial cells (hAECs) from 4 healthy donors (Supplementary Table 1) were seeded in the Airway Lung-Chips on a human placenta collagen IV-coated 3-µm pore polyester membrane at a density of 3 × 10^6^ cells/mL and left to attach for 2h. Five days post seeding, air liquid interface (ALI) was introduced and Airway Lung-Chips were perfused basally at 60µL/h for 3 weeks until full differentiation. Epithelium integrity, ciliation, and apical mucus secretion were monitored for quality control. Upon full epithelial differentation, hMVECs or HUVECs were seeded onto the opposite side of the membrane, in the vascular channel at a density of 1 × 10^7^ cells/mL and cultured under flow for 3-4 days in endothelial growth media. Rhinovirus strain 16 (HRV-16) (ATCC® VR-283) was inoculated into the epithelial channel of fully differentiated Airway Lung-Chips and incubated for 3h at 33°C while control chips received DMEM only. Following incubation, the epithelial channel of each chip was gently rinsed 5 x with DMEM. After the final wash, and every 24h thereafter, the epithelial surface was washed with 50 µL of DMEM to determine virus growth kinetics and the basolateral effluent was collected for cytokine/chemokine responses. HRV16 titers were determined by infecting H1-HeLa cells (CRL-1958) with serially diluted virus samples, recording the number of wells positive for viral cytopathic effect (CPE) at each dilution after 5 days post infection and calculating the TCID_50_/mL. A Th2 microenvironment was induced by perfusing the vascular channel of the Airway Lung-Chips with 100 ng/mL of IL-13 (Peprotech, USA) for 7 days. High-content imaging was performed in live and fixed tissues and followed by quantitative analysis to extract structural and kinematic endpoints, including ciliary beat frequency and neutrophil recruitment dynamics. Anti-dsRNA antibody J2 (mabJ2, scicons, CZR) was utilized to detect replicating intracellular HRV16 by immunofluorescence staining.

## Results

Here, we describe the development of a human Airway Lung-Chip suitable for the study of viral-induced asthma exacerbation and the effect of immunomodulatory compounds on immune cell dynamics (**Supplementary Fig. 1A**).

### Adapting the Airway Lung-Chip design to enable immune cell transmigration

We adapted our recently developed micro-engineered “small-airway-on-a-chip” model ^10^ by increasing the membrane pore size (3.0 µm vs 0.4 µm) to allow immune cells to transmigrate from the vascular microchannel to the epithelial lumen (**Supplementary Fig. 1B**). As substrate pore size can greatly influence tissue differentiation^14^, we validated the new chip design for airway tissue culture. hAECs were seeded in the top channel and human primary endothelial cells were seeded in the bottom channel to create a tissue-tissue interface. We confirmed establishment of a well-differentiated mucociliary bronchiolar epithelium by the presence of robust ZO-1-containing tight junctions, characteristic epithelial cobblestone-like morphology, mucus producing goblet cells, and dense coverage with ciliated cells (**Supplementary Fig. 1C-E)**. Formation of an intact microvascular endothelium was validated by staining for endothelial junctional protein VE-Cadherin (**Supplementary Fig. 1F)**. Mucociliary function was assessed by high-speed video microscopy and quantitative analysis^15^. We measured a ciliary beat frequency of 16.35 (± 2.6) Hz (**Supplementary Fig. 1G-H, Supplementary movie 1**). Microbead trajectories revealed locally unidirectional mucociliary transport (**Supplementary Fig. 1I**) with a typical velocity of ∼ 100 µm/sec, which is similar to mucociliary transport velocities in intact human airways ^16^. In sum, the Airway Lung-Chip was functionally augmented with a 3-µm pore membrane to enable immune cell transmigration, and we validated that it supports the viability and function of normal human bronchiolar tissue.

### HRV16 readily infects the Airway Lung-Chip and induces cytopathic effects

We next investigated HRV infection of the Airway Lung-Chip (**Fig. 1A)**. We infected the chip apically with HRV16, a rhinovirus serotype commonly used in clinical challenge studies, and analyzed viral infectivity and replication, cell tropism and induced cytopathic effects (CPE). Infection a multiplicity of infection (MOI) of 1 resulted in a rapid apical release of infectious virus particles that peaked between 24 and 48 hours post infection (hpi) followed by a steady decline of released virions until no virus was detected 144 hpi (**Fig. 1B**). This timeframe is consistent with viral shedding and symptom score profiles of individuals experimentally infected with HRV ^17–19^.

**Fig 1.**
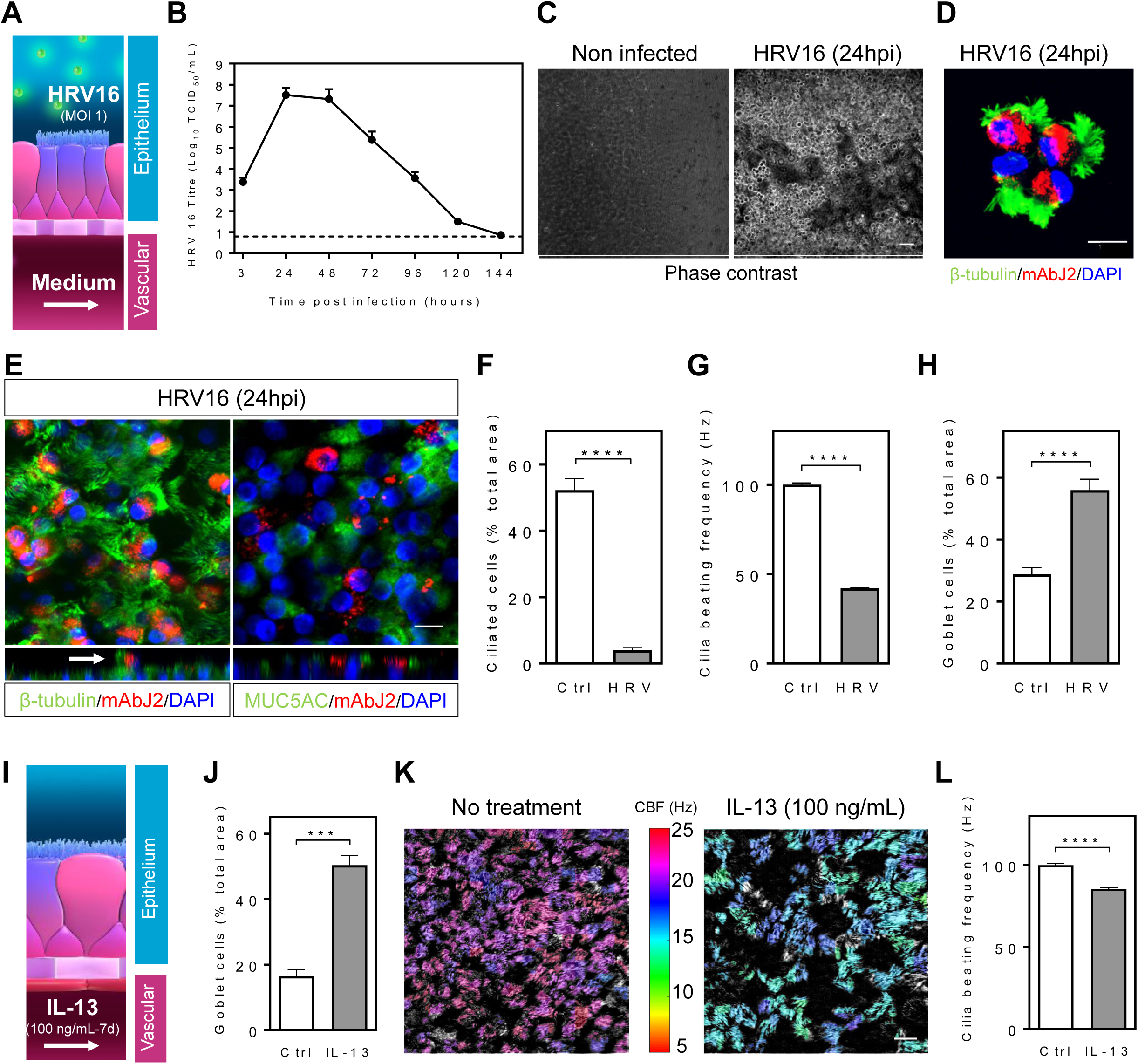
Modeling HRV infection and Th2 asthma phenotype on the Airway Lung-Chip. (**A**) HRV-16 was apically delivered to the chip at an MOI∼1. (**B**) Viral growth kinetic of HRV-16 following infection. HRV16 was titrated in apical washes and data represent mean ± SD log10 TCID_50_/ml from 3 different donors with two to five biological replicates per donor. The dashed line indicate the limit of detection of the titration assay. (**C**) Phase contrast micrographs of differentiated airway epithelium cultured in the airway chip for 3 weeks at air liquid interface infected with HRV16 for 24h (right) or non infected (left) showing epithelial cells sloughed off infected cultures while non infected epithelium remains intact. Scale bar, 50 µm. (**D**) IF image of apical washes from cultures infected with HRV16 for 24h showing detached cells stained for β-tubulin (green), mAbJ2 (red) and DAPI (blue). Scale bar, 10 µm. (**E**) IF image of the differentiated mucociliary epithelium on-chip infected with HRV16 for 24h and stained for β-tubulin (left) or MUC5AC (right) (green) and mAbJ2 (red). Orthogonal views show a ciliated infected cells protruding from the epithelial lining (white arrow). Scale bar, 20 µm. (**F**) Total culture area covered by ciliated cells in non infected chips and 6 days post HRV16 infection. (**G**) Cilia beating frequency in non infected chips and 72 hpi. (**H**) Total culture area covered by goblet cells in non infected control and HRV16 infected chips 6 dpi. (**I**) A Th2 asthma environment was generated through vascular perfusion of IL-13 (100 ng/mL) over 7 days. (**J**) Total culture area covered by goblet cells in absence or presence of IL-13 (100 ng/mL) for 7 days. (**K**) Heat map of cilia beating frequency (CBF) on-chip generated from a high speed recording of cilia beating in a representative field of view in absence or presence of IL-13 (100 ng/mL). Scale shows color coded CBF values in Hz. Scale bar, 20 μm. (**L**) Graph showing ciliary beat frequency in absence or presence of IL-13 (100 ng/mL). Data are from chips with cells from three healthy donors, with one or two biological replicates (chips) per donor. Data represent mean ±SEM and significance was determined by unpaired Student’s; ****P < 0.0001.

HRV16 readily infected the intact epithelium and induced apical cell sloughing by 24 hpi (**Fig. 1C**). Real time imaging and immunofluorescence (IF) characterization identified that the majority of detached cells were infected multiciliated cells (**Fig. 1D, Supplementary movie 2**), similar to what has been observed in respiratory tissue biopsies of human volunteers experimentally infected with HRV^20,21^. Many of the detached cells were TUNEL-positive, indicating apoptosis (**Supplementary Fig. 2A**). IF imaging of infected chips revealed that HRV16 infection was restricted to ciliated cells and induced the protrusion and extrusion of infected cells from the epithelial lining, while mucus producing cells remained non-infected (**Fig.1E, Supplementary movie 2**). By day 6 post infection, few ciliated cells remained on the epithelial surface (13-fold decrease in cilia coverage; p<0.001) (**Fig. 1F, Supplementary Fig. 2B**) and they displayed significantly shorter cilia (**Supplementary Fig 2C-D**). Additionally, consistent with HRV16 ciliated cell tropism, we observed a 60% reduction in ciliary beat frequency following infection (p<0.001) (**Fig. 1G** and **Supplementary movie 3**). The acute damage to ciliated cells was accompanied by a 2-fold increase in the area covered by MUC5AC positive cells by day 6 post infection compared to non-infected chips (p<0.001) (**Fig. 1H, Supplementary Fig. 2E**), in line with clinical observations^22^.

### Modeling viral-induced asthma exacerbation on-chip

We used interleukin-13 (IL-13) stimulation to induce features of type 2 inflammation in asthma (**Fig. 1I**). IL-13 plays a central role in allergic asthma and, together with IL-5, is an important mediator of viral-induced exacerbations in asthma ^23^. When we perfused the vascular channel of the Airway Lung-Chip with IL-13 (100 ng/mL) for 7 days preceding virus infection we detected a significant increase in the number of goblet cells (rising from 16.5% to 50.3% total area; p<0.01) (**Fig. 1K, Supplementary Fig. 3**) and a decrease in cilia beating frequency (14.4% reduction; p<0.001) (**Fig. 1K,L and Supplementary movie 4**), which is consistent with observations made in airway mucosa of asthmatic patients ^24,25^.

To recapitulate HRV-induced asthma exacerbation, we combined IL-13 treatment with HRV16 infection to induce exacerbation (**Fig. 2A**). Infection of the IL-13 stimulated Airway Lung-Chip with HRV16 did not result in significant changes in viral loads in apical washes nor in the duration of infection when compared with non-stimulated chips (**Fig. 2B**). This finding is consistent with recent studies in asthmatics experimentally infected with HRV16 ^26–28^, although another study found a higher viral load in asthmatics on day 3 of infection^29^.

**Figure 2.**
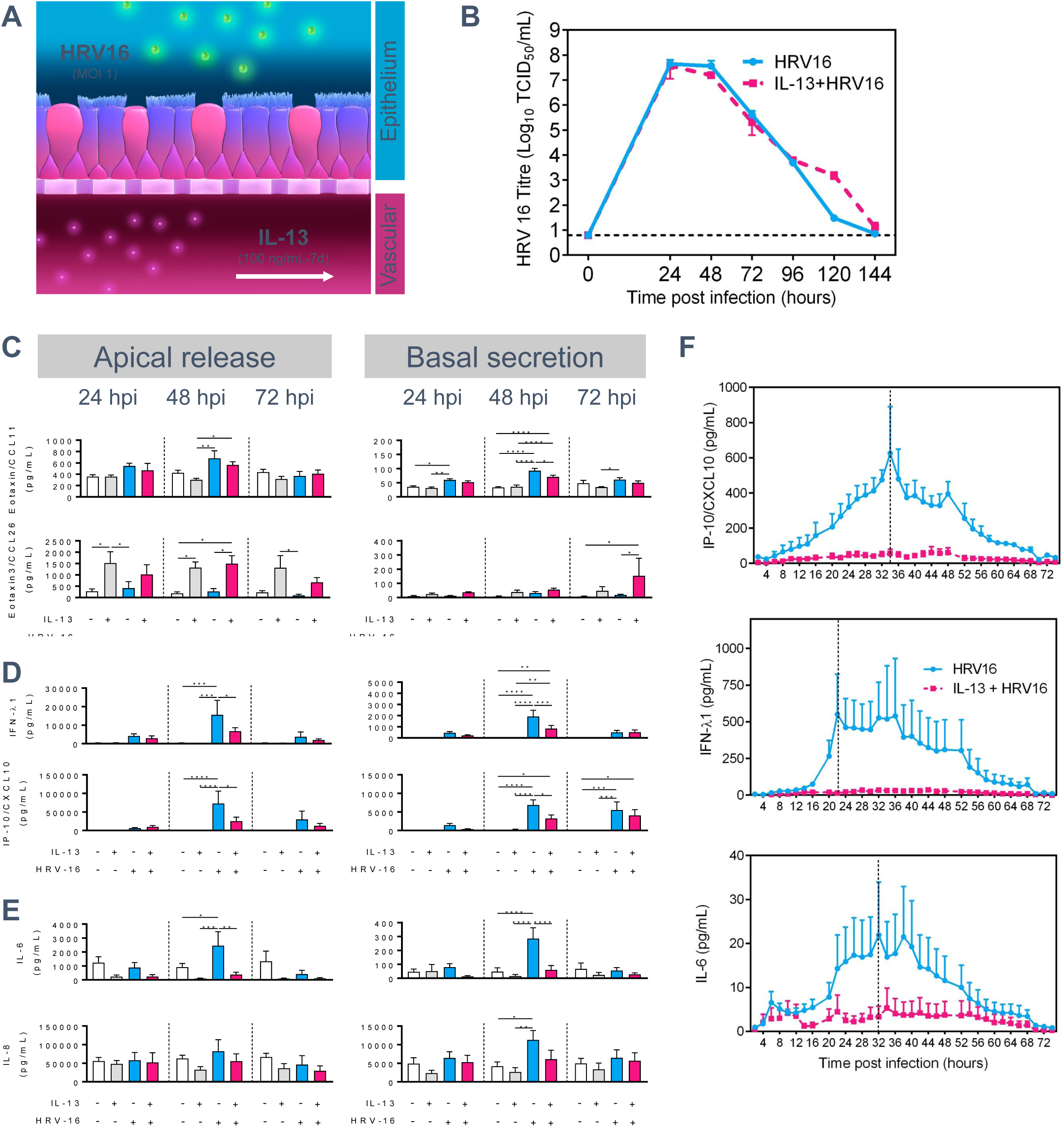
Modeling HRV-induced asthma exacerbation on chip and measuring inflammatory responses. (**A**) The severe asthma chip was generated by inducing features of Th2 asthma through vascular perfusion of IL-13 (100 ng/mL) and exacerbation through HRV16 epithelial infection (MOI 1). (**B**) Viral growth kinetic of HRV-16 following infection of the airway chip at an MOI∼1 in the absence or presence of IL-13. HRV-16 was titrated in apical washes and data represent mean ± SEM log10 TCID_50_/ml from 3 different donors with two to five biological replicates per donor. **(C-E)** The graphs show the effect of on polarized apical and basal release of CC chemokines (**C**), interferon and interferon-induced chemokines (**D**), and pro-inflammatory interleukins (E) by the combination of IL-13 treatment and HRV-16 infection in the exacerbated asthma chip at 24h, 48h and 72h post infection. (**F**) The graphs show a high resolution, kinetic profiles of insert IFN-λ1, IP-10/CXCL10and IL-6 response following HRV16 infection, with and without IL-13 treatment over 72 h. Data represent mean ±SEM of cells from three to four different donors, with one or two biological replicates (chips) per donor. Significance was determined by multiple comparison two-way ANOVA followed by Tukey’s post-hoc correction; *P < 0.05, **P < 0.01, ***P < 0.001, ****P < 0.0001.

Type 2 inflammation has been suggested to increase asthmatics susceptibility to exacerbations through airway remodeling and modulation of the antiviral response ^30^. To gain insight into the effect of IL-13 on HRV16-induced inflammatory responses we first measured the levels of pro-inflammatory cytokines and chemokines released apically and basolaterally at 24, 48 and 72 h post HRV16 infection, in the presence or absence of IL-13 (**Fig.2C-E and Supplementary Fig. 4**). Apical and basolateral secretion of CCL11 (Eotaxin-1), a potent eosinophil chemoattractant and marker of Th2 asthma in human ^31^, was increased following HRV16 infection (**Fig. 2C**), consistent with literature^32^. This was not significantly altered by IL-13 treatment. In contrast and consistent with literature^33^, apical secretion of CCL26 (Eotaxin-3), another chemokine associated with eosinophil recruitment and persistence in patients with asthma^34^, was increased at all time points following stimulation with IL-13 alone, while HRV16 infection had no effect on CCL26 release. Analysis of the interferon (IFN) type III responses revealed that apical and basal release of IFN-λ1 at 48 hpi, and of the IFN-induced lymphocyte chemoattractant CXCL10 at 48 and 72 hpi were increased following HRV16 infection (**Fig. 2D**). IL-13 treatment significantly reduced the measured IFN response in HRV16 infected chips. This is consistent with other in vitro studies showing that IL-13 impairs IFN-λ production^35^, and with clinical reports intimating that individuals with asthma or cells from asthmatics infected with HRV have a deficient IFN response ^36–39^, although this remains controversial^40^. Basal secretion of IL-8, a potent attractant of neutrophils^41^, was increased 48 h post HRV16 infection, with little impact of IL-13 treatment (**Fig. 2E**). In contrast, we found that induction of IL-6 secretion following HRV16 infection was greatly reduced in IL-13 treated samples at 48 h post infection (**Fig. 2E)**. Notably, we did not detect any IFN-β secretion at any time point with and without IL-13 stimulation.

To elucidate the kinetics of inflammatory mediator release during viral-induced asthma exacerbations and thereby gain additional insight into the pathophysiological process^42,43^, we exploited the microperfusion of the Airway Lung-Chip for high resolution temporal profiling of IFN-λ1, CXCL10 and IL-6 secretion. By collecting basal medium outflow every 2h for 74h, we revealed the temporal dynamics of cytokine secretion after HRV infection (**Fig. 2F**). In untreated chips, both IFN-λ1 and IL-6 increased quickly and reached maximal, or near maximal, values at approximately 22 h post HRV16 infection. CXCL10 rose more slowly and peaked at 34 h post infection, in line with direct interferon-dependent regulation of CXCL10. In contrast, the IL-13 treated chips exhibited markedly suppressed secretion profiles. The high-resolution profiling plots reveal this difference in the first 24h, whereas it remains masked in the accumulated medium collected after 24 hpi (**Fig. 2D,E**). This demonstrates that high resolution profiling is a sensitive method for detecting onset and dynamics of cytokines release, determining sensitive time windows, and identifying the best sampling strategy.

We also investigated the effect of the IL-13 stimulation of human endothelial cells, which contributes to leukocyte recruitment in the pathogenesis of asthma^44^. Within 48h of the IL-13 treatment, we observed progressive endothelium morphological changes leading to the formation of cell aggregates throughout the vascular wall (**Supplementary Fig. 5A**) accompanied by a doubling of endothelial cell density (p < 0.0001) (**Supplementary Fig. 5B**). Further analysis revealed increased gene expression of P-Selectin (p = 0.036) and E-Selectin (p = 0.06), two adhesion molecules involved in inflammatory cell recruitment, compared to control (**Supplementary Fig. 5C, Supplementary Table 2**). IF staining of intercellular adhesion molecule 1 (ICAM-1) and vascular cell adhesion protein 1 (VCAM-1) was also increased in IL-13 treated cultures (**Supplementary Fig. 5E)**, indicating that IL-13 promotes activation of the lung microvascular endothelium. Indeed, bronchial biopsies from subjects with asthma revealed increased adhesion molecule expression in endothelial cells^45^.Although we cannot rule out that the IL-13 treatment stimulated the epithelial cells to release additional chemokines, our findings align with previous studies on the effects of IL-13 on human endothelial cells^46^.

### Pharmacological modulation of neutrophil recruitment and transmigration by a CXCR2 antagonist on-chip

Airway neutrophilia is frequently observed in acute exacerbations of asthma^13,47,48^, and is associated with asthma severity^49^. HRV infections in asthmatics may also promote selective recruitment of neutrophils^50^. Since neutrophils can contribute to asthma exacerbation through the release of tissue-damaging compounds^51^, neutrophil recruitment constitutes a potential drug target. We sought to show feasibility of probing the real-time dynamics of neutrophil recruitment on the Airway Lung-Chip, where the endothelial vascular channel supports the live observation of circulating and transmigrating neutrophils. In a proof-of-concept study, we investigated neutrophil recruitment in exacerbated conditions and in response to a CXCR2 antagonist, which blocks neutrophil sensing of IL-8^41^. Here, we perfused freshly isolated human neutrophils stained with a live dye through the vascular channel of exacerbated asthma chips under physiological flow (i.e., with fluid shear stresses of ca. 1 dyn/cm^2 52^, an important factor for physiological neutrophil recruitment^53^). We observed neutrophil adhesion and crawling at the surface of the microvascular endothelium, rapidly followed by neutrophil transendothelial migration through a combination of transcellular and paracellular migratory events (**Fig. 3A, B**). Treatment with either IL-13 or HRV16 significantly increased neutrophil adhesion, and combined treatment almost doubled recruitment compared to each condition alone (**Fig. 3C, D**). Pre-treatment of neutrophils with 10 µM MK-7123, a CXCR2 antagonist, at 24 hpi significantly reduced neutrophil adhesion to the endothelium in all treatment groups (**Fig. 3C, D**), and also reduced neutrophil crawling velocity (**Fig. 3E, F and Supplementary movies 5 and 6**). Real time imaging revealed that many endothelium bound neutrophils transmigrated from the vascular channel through the 3-µm pores of the membrane (which frequently merge into even larger pores in track-etched membranes) into the epithelium channel where they adhered to the epithelial surface (**Fig. 3G and Supplementary movie 7**) and treatment with MK-7123 reduced the proportion of transmigrating neutrophils (**Fig. 3H**). Furthermore, characterization of fully migrated neutrophils revealed expression of myeloperoxidase (**Fig. 3I**), a marker of neutrophil activation found in lungs of asthmatics ^54^ that contributes to inflammation *in vivo* ^55^. Thus, our on-chip disease model of asthma exacerbation not only enabled efficacy testing of an immunomodulatory compound but also elucidated the multi-step dynamics of neutrophil recruitment in a human-relevant context.

**Fig 3.**
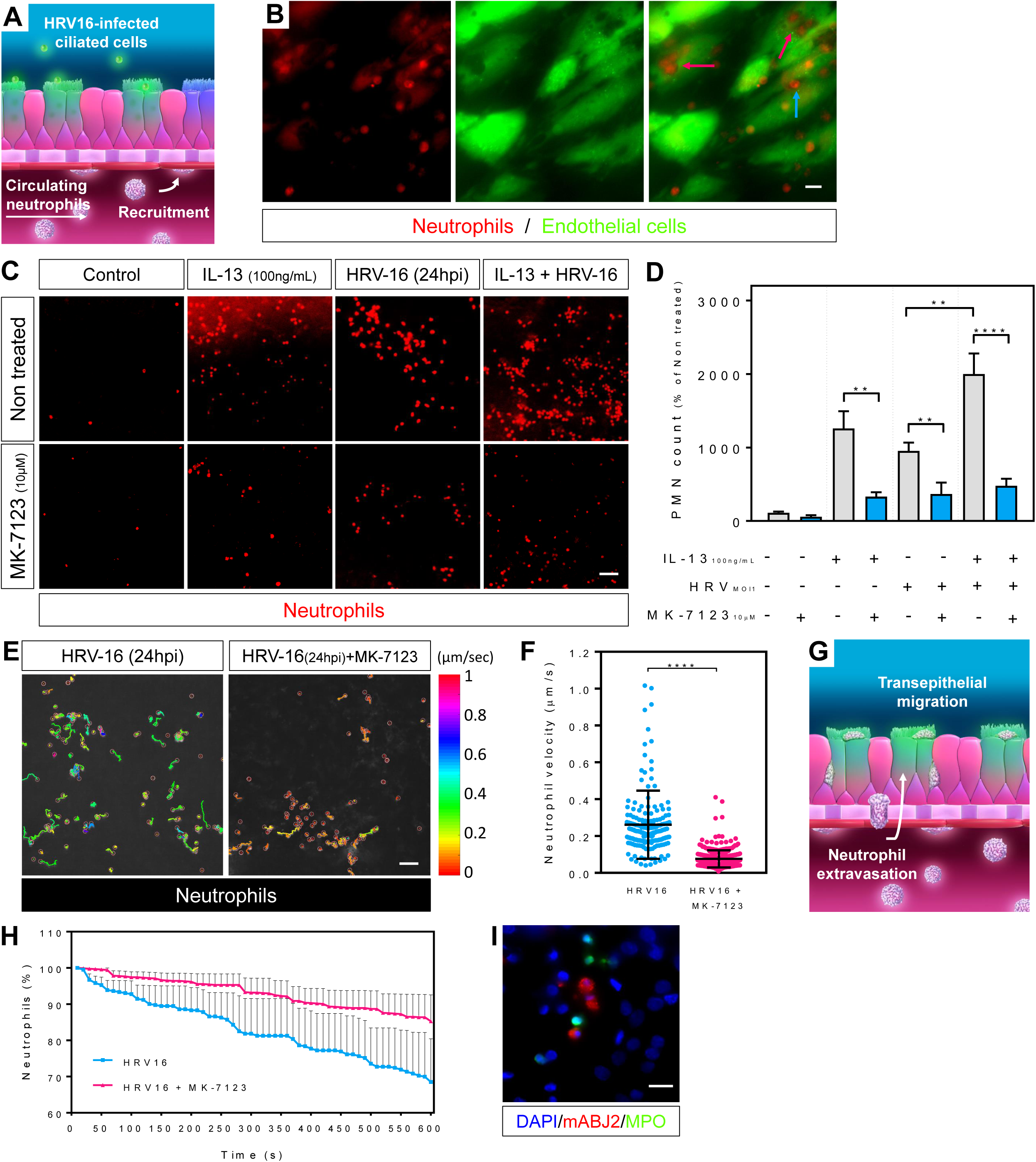
Assessing and modulating IL-13- and HRV-induced neutrophil recruitment and infiltration on chip. (**A**) Schematic diagram of the severe asthma chip depicting the neutrophil recruitment assay. (**B**) Micrographs showing paracellular and transcellular transmigration of neutrophil live-stained (red) through GFP-expressing endothelial cells (green). Obvious transcellular diapedesis events are indicated by the pink arrows while the blue arrow points at a paracellular migration. Scale bar, 10 µm. (**C**) Fluorescence microscopic imaging of chips perfused through the microvascular channel under physiological conditions (shear stress, 1 dyn/cm^2^) for 5 min with freshly isolated CellTracker red-labeled human neutrophils showing recruitment to the endothelium stimulated or not with a combination of IL-13, HRV-16 and a CXCR2 antagonist (MK-7123 - 10µM). Scale bar, 50 µm. (**D**) Quantification of neutrophil recruitment from (**C**). (**E**) Schematic diagram of neutrophil diapedesis taking place in the severe asthma chip. (**F**) Fluorescence imaging tracking neutrophil velocity at the endothelial surface in HRV-16 infected chips, in absence (left) or presence (right) of the CXCR2 antagonist. Scale shows color coded velocity values in µm/sec; Scale bar, 50 µm. (**G**) Quantification of neutrophil velocity in HRV16 infected chips with or without MK-7123 (10µM). (**H**) The graph shows the disappearance of neutrophils (expressed in % of starting count) due to transmigration from the vascular wall to the epithelial lumen in HRV16 infected chips in absence or presence of MK-7123 (10µM). (**I**) Immunofluorescence micrograph of an HRV16 infected chip showing human neutrophil stained for myeloperoxidase (green) that have migrated to the epithelial lumen among HRV16 infected epithelial cells (red). Scale bar, 20 µm. Data represent mean ± SEM from three different donors with two biological replicates (chips) per donor.

## Discussion

We still incompletely understand why asthmatics are at higher risk of developing life-threatening exacerbation following rhinovirus infection. Previous *in vitro* studies have suggested that prior damage to the asthmatic airway epithelium facilitates HRV infection by exposing more permissive, ICAM-1-expressing basal cells, explaining the susceptibility of asthmatic airways to infection ^56,57^. The corollary is the perception that intact epithelial layers of healthy subjects are somewhat resistant to HRV infection^58^. Contrasting with this view, our findings demonstrate that HRV can readily infect the intact well-differentiated mucociliary epithelium and induce significant airway remodeling leading to almost complete loss of ciliated cells and goblet cell hyperplasia by day 7 post infection. Our observations are consistent with studies showing induction of mucin production by rhinovirus infection of human bronchial epithelia in vitro and in vivo ^22,59^, the induction of goblet cell metaplasia by rhinovirus-mediated foxa3 activation^60^, high viral load and cell shedding the airways of individuals infected with HRV^20,26–28^, as well as HRV-induced CPEs^61,62^ and apoptotic events^63^ in *in vitro* human bronchial epithelia. This suggests that RV infections have considerable potential for cytopathic and remodeling effects in the airways, and that HRV-induced CPE might contribute to exacerbations in asthmatic individuals by directly damaging mucociliary clearance and increasing mucus secretion.

Several groups have investigated the effect of IL-13-induced mucous metaplasia on the susceptibility of human airway epithelium to HRV infection and come to discordant conclusions, such as increased versus reduced susceptibility to HRV infection ^64 65^. In contrast, we did not detect any significant difference in viral load or infection persistence in IL-13 treated versus non-treated chips. Donor to donor variability and differences in experimental design might explain the different outcomes. Clinical studies are also in disagreement, with some reporting higher viral loads in asthmatics^29^ while others align with our results in showing no difference in viral titer of asthmatics compared to non-asthmatics ^26–28,66^. In sum, there are clearly many remaining unknowns in HRV-induced asthma exacerbation, and our findings show that the Airway Lung-Chip is a valuable new tool for their investigation.

We found that HRV-induced type III IFN response was ablated in a type 2 microenvironment, which is consistent with studies showing impaired IFN responses in cells from asthmatic individuals challenged with HRV ^36,67^. However, several recent *in vitro* and *in vivo* studies contradict these results ^26,28,40,68,69^, suggesting that an impaired IFN responses may not be the only factor contributing to exacerbation. Asthma phenotype heterogeneity and differences in the study designs may further confound the picture. An impaired IFN response might be specific to asthmatics with a Th2-dominant inflammatory profile ^23,36^, in line with the downregulation of IFN-λ1 by a Th2 microenvironment on our chip.

Effective orchestration and resolution of the immune response rely on both magnitude and kinetics of secreted cytokines. IL-6 is a prime example as its pro- or anti-inflammatory nature depends on secretion timing ^43^. Here, we have leveraged the microscale perfusion of the Airway Lung-Chip to conduct high resolution temporal profiling of secreted cytokines. We found that the IL-6 secretory profile was markedly disrupted by IL-13 stimulation. This could be due to the altered epithelial architecture and cell type composition of the asthmatic microenvironment^70^, which may alter cytokine responses^71,72^. The central role of IL-6 in infection and inflammation has been extensively described ^73^ and though it is usually considered a pro-inflammatory mediator in asthma exacerbations, its protective role in the lung was highlighted in several studies ^42 74 75 76^, which together with our results may suggest a more nuanced role for IL-6 in exacerbation pathogenesis.

We also demonstrated the utility of the Airway Lung-Chip for testing the efficacy of immunomodulatory compounds on neutrophil recruitment. Though the first generation of CXCR2 inhibitors has failed to deliver in the clinic, it is now appreciated that neutrophils are extremely heterogeneous and display different responses depending on subtype and environmental factors^77^. Because of this, more than ever new platforms like the Airway Lung-Chip are needed to enable the phenotypic and functional profiling of neutrophils in the context of disease and drug treatment.

In summary, we developed a new on-chip human disease model of viral-induced asthma exacerbations that recapitulates hallmarks of rhinovirus-induced responses of the asthmatic airway. Owing to its unique fluidic design and accessibility to optical imaging, the Airway Lung-Chip also enables the high-resolution profiling of cytokine release, phenotypic changes and dynamic immune cell responses to immunomodulatory therapies in a human-relevant context, thus adding a powerful pre-clinical platform for probing complex disease mechanisms and developing new treatments for asthma exacerbations.

## Acknowledgments

We thank Dr. Anne van der Does and Dr. Gurpreet Brar for the critical review of the manuscript and Brett Clair for his assistance with graphic design.

---

**Supplementary figure 1.**
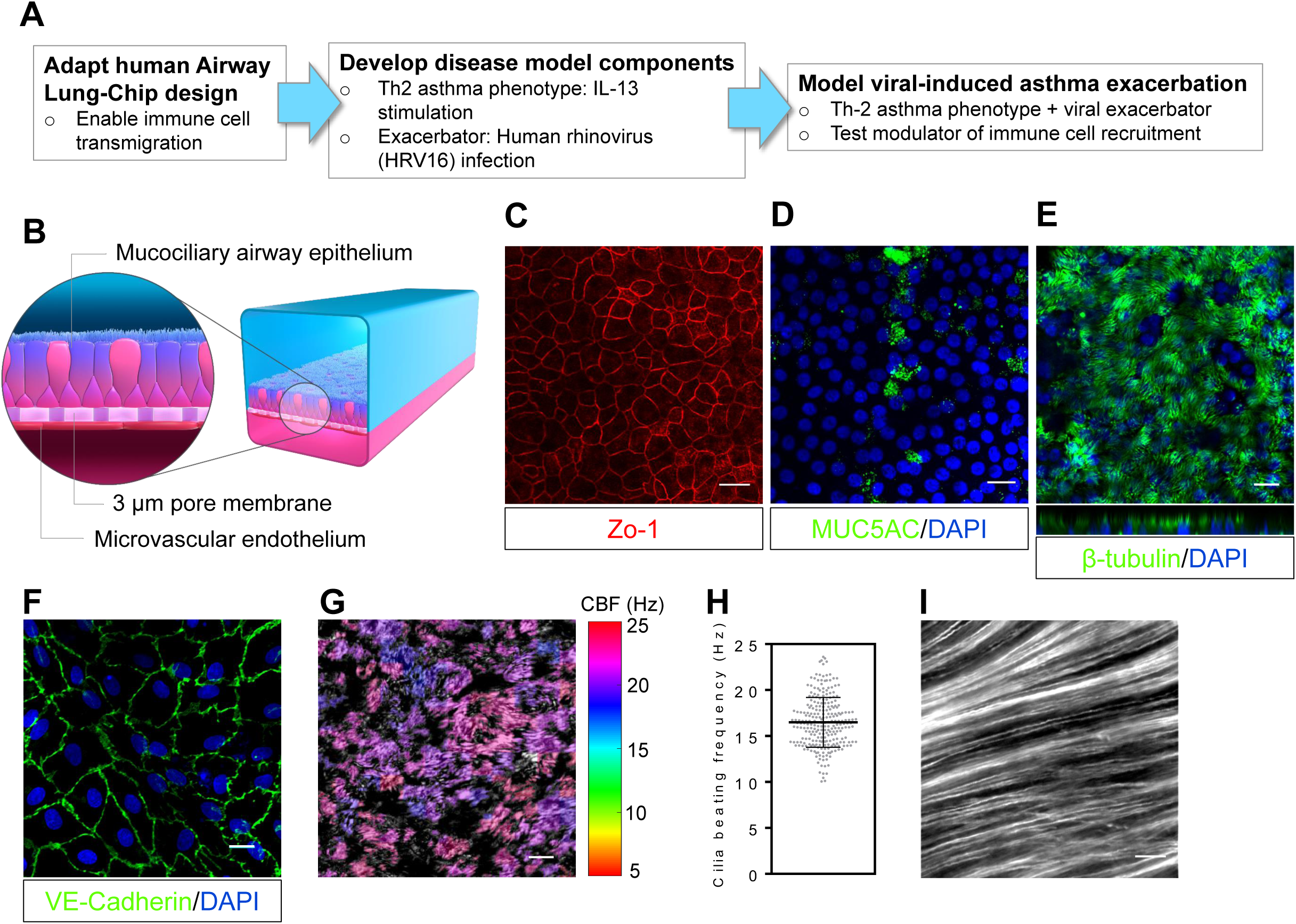
The human Airway chip. **A**) Overview of approach taken in this study to model asthma exacerbation and demonstrate feasibility of compound testing. (**B**) Functionally augmented Airway Lung-Chip design has 3-µm pore membrane for enabling neutrophil transmigration between the endothelial and epithelial channel. (**C**) The differentiated airway epithelium on the top channel exhibits continuous tight junctional connections on-chip, as demonstrated by Zo-1 staining (red). Scale bar, 20 μm. (**D-E**) The well-differentiated human airway epithelium generated on-chip also contains goblet cells stained for MUC5AC (green) and demonstrates extensive coverage of ciliated cells labelled for β-tubulin (green). Scale bar, 20 μm. (**F**) The human (lung blood microvascular) endothelial monolayer formed in the bottom channel of the chip features continuous adherens junctions between adjacent cells, as indicated by VE-Cadherin staining (green). Scale bar, 10 μm. All images are representative of two to five independent experiments performed with cells from three different donors. (**G**) Graphic representation of cilia beating frequency (CBF) on-chip generated from a high speed recording of cilia beating in a representative field of view. Scale shows color coded CBF values in Hz. Scale bar, 20 μm. (**H**) The graph shows the cilia beating frequency measured in 5-10 random on-chip fields of view and each dot represents regions of ciliary beating. Data have been recorded in cells from three different donors and are presented as mean ± SD. (**I**) Time lapse composite image of a 5 sec recording of 1-µm fluorescent microbeads suspended in PBS and introduced in the top channel. The path lines reveal the mucociliary transport of the differentiated hAECs. Scale bar, 20 µm.

**Supplementary figure 2.**
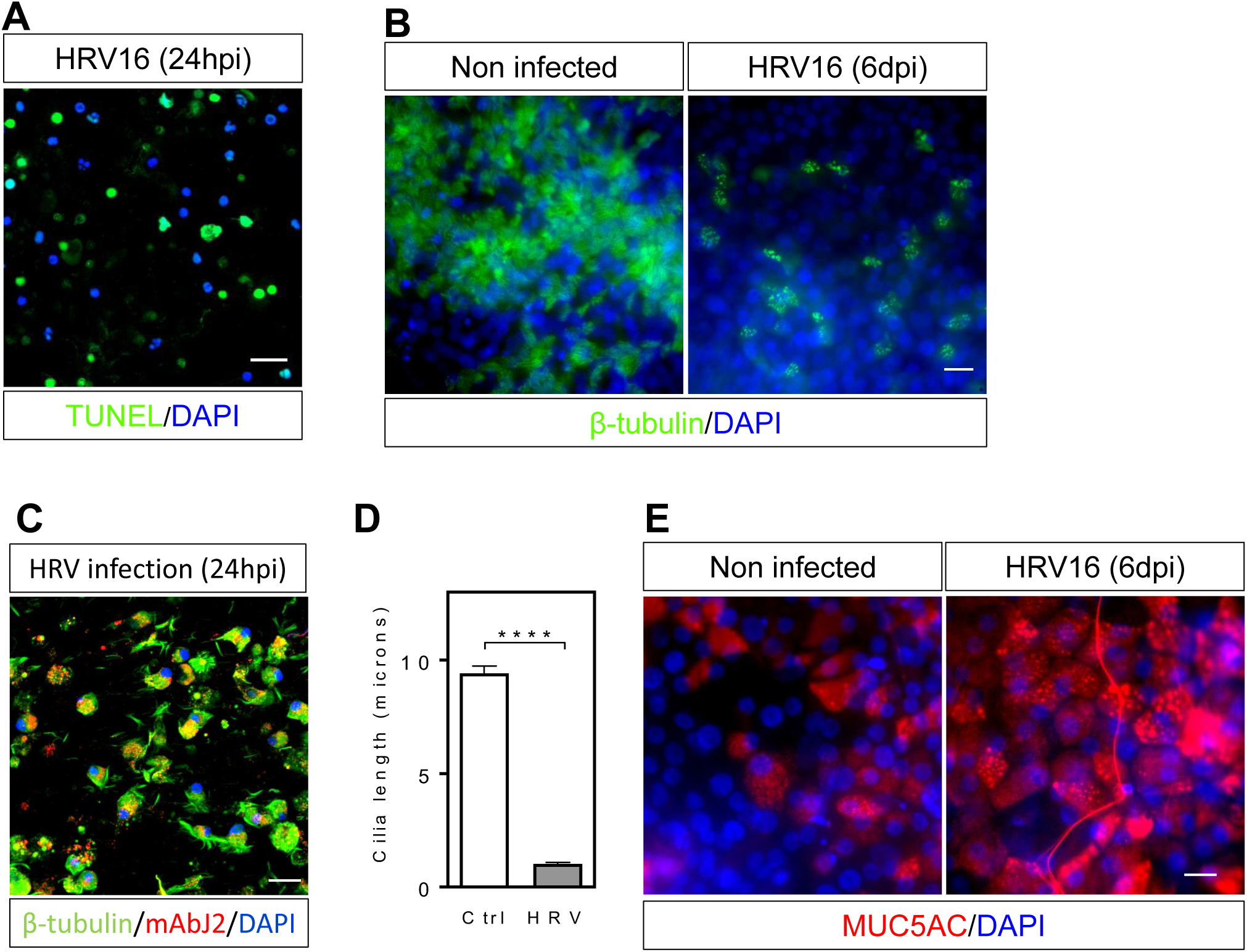
HRV infection causes cytopathic effects and epithelial remodeling. (**A**) Fluorescence confocal micrograph of detached epithelial cells stained using the TUNEL assay showing nuclei of apoptotic (green) and non apoptotic (blue) cells. Scale bar, 20 µm. (**B**) Immunofluorescence images of the differentiated epithelium non infected (left) or fixed at 6 days post HRV16 infection (right) and stained for β-tubulin (green) and DAPI (blue). Scale bar, 20 µm. (**D**) Confocal image of the cells collected in an apical wash 24 h following HRV16 infection of an airway chip showing the extend of ciliated cells destruction. Note the presence of individual cilia (green) in the apical wash. Scale bar, 20 µm. (**D**) Quantification of the length of cilia on multiciliated cells 6 d post HRV16 infection. Data represent mean ± SEM; Significance was determined by unpaired Student’s t-test; ****P < 0.0001. (**E**) Immunofluorescence images of the differentiated epithelium non infected (left) or 6 days post HRV16 infection (right) and stained for MUC5AC (red) and DAPI (blue). Scale bar, 20 µm.

**Supplementary Figure 3.**
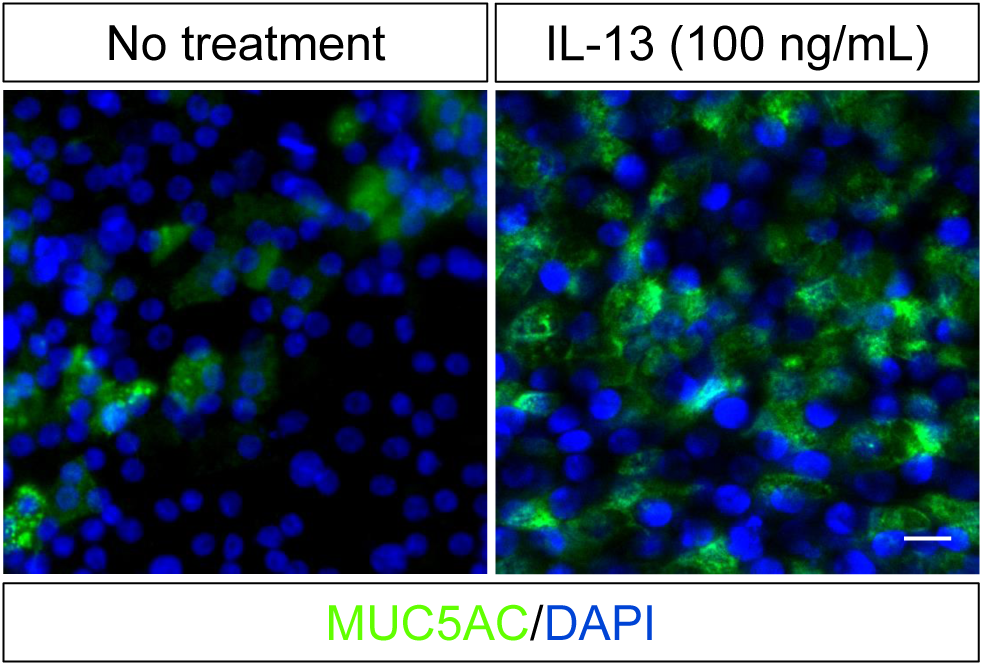
IL-13 induces goblet cell hyperplasia. Immunofluorescence confocal views of differentiated airway epithelium cultured at air-liquid interface for 3 weeks on-chip in the absence (left) or presence (right) of IL-13 (100 ng/mL) for 7 days showing epithelium stained for the goblet cell marker MUC5AC (green) and nuclei (blue). Scale bar, 20 μm (applies to both views); images are representative of two independent experiments performed on cells from three different donors.

**Supplementary figure 4.**
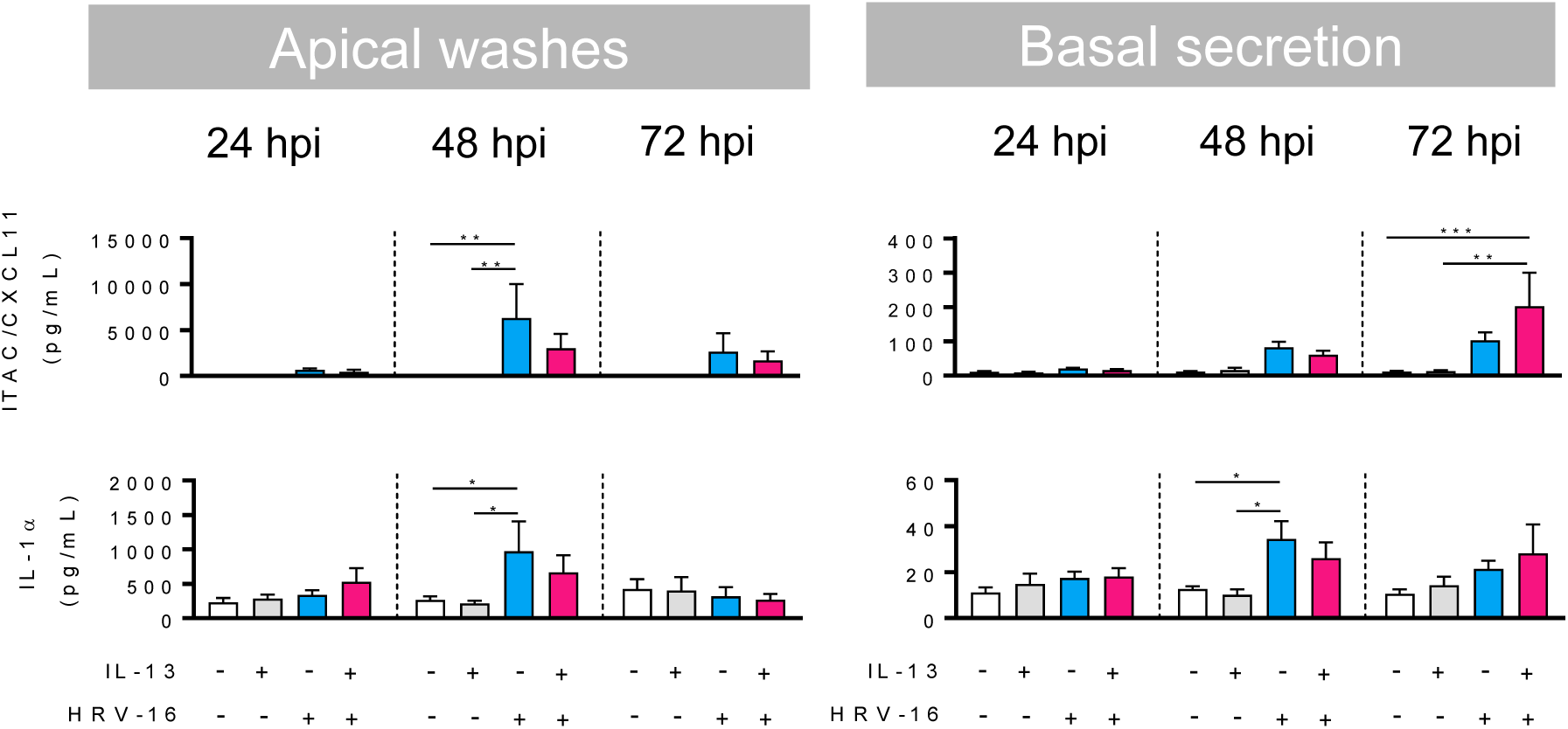
The graphs show the effect on polarized apical and basal release of CXCL11 and IL-1α by the combination of IL-13 treatment and HRV-16 infection in the severe asthma-on-a-chip at 24h, 48h and 72h post infection. Data represent mean ±SEM of cells from three to four different donors, with one or two biological replicates (chips) per donor. Significance was determined by multiple comparison two-way ANOVA followed by Tukey’s post-hoc correction; *P < 0.05, **P < 0.01, ***P < 0.001.

**Supplementary figure 5.**
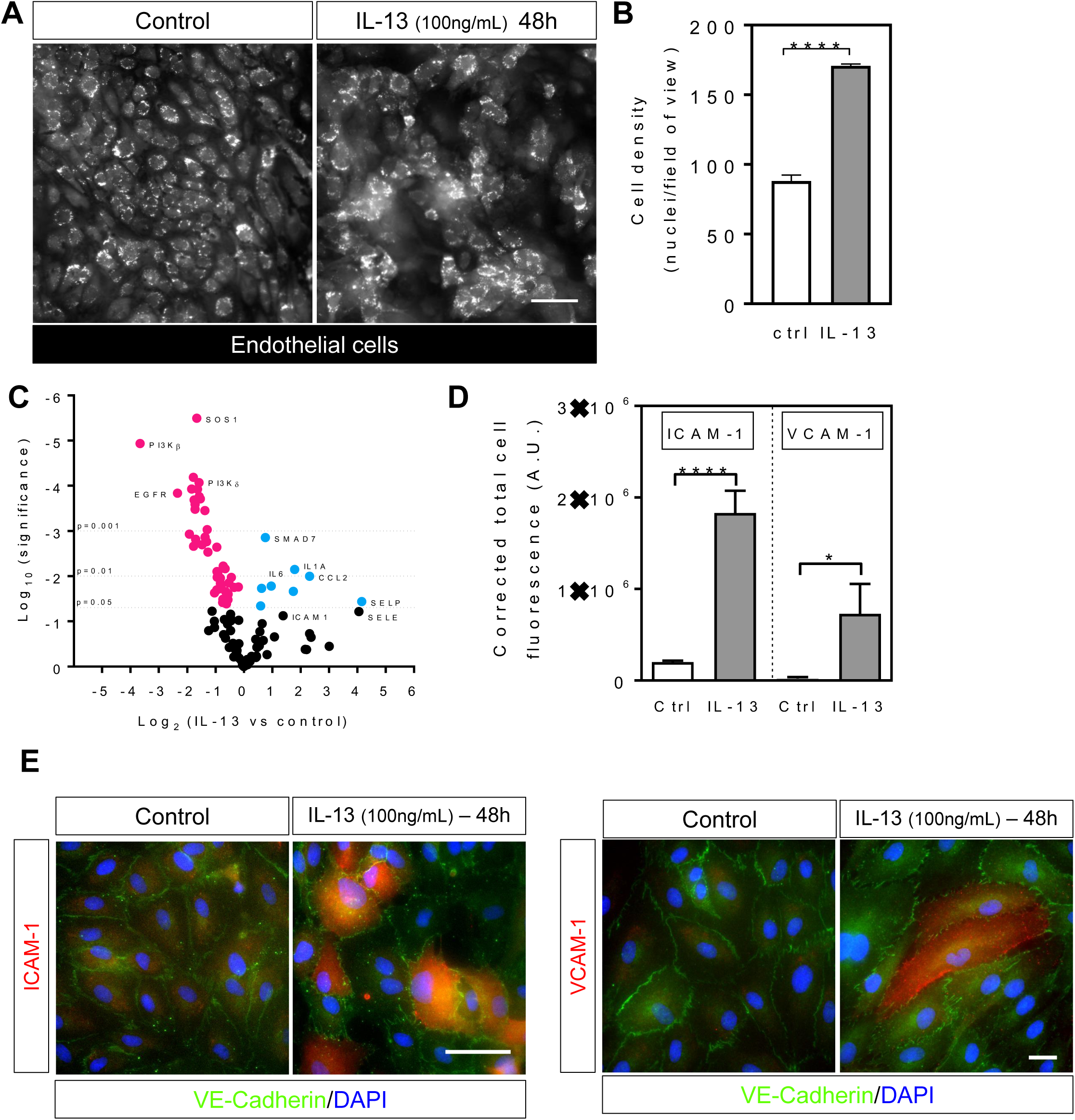
IL-13 induces aggregation and activation of endothelial cells. (**A**) Fluorescence micrographs of GFP-expressing endothelial cells co-cultured with well differentiated airway epithelial cells on chip, in absence (left) or presence (right) of IL-13 (100ng/mL) for 48h. Scale bar, 50 µm. (**B**) Quantification of endothelial cell density in absence or presence of IL-13 (100 ng/mL) for 48h. (**C**) Volcano plot showing gene expression of IL-13 (100ng/mL) treated endothelial cells compared to untreated group. Data were analyzed using a two-stage step-up method of Benjamini, Krieger and Yekutieli. (**D**) Effect of IL-13 (100ng/mL) on endothelial cell adhesion molecules ICAM-1 and VCAM-1 expression, as measured using fluorescence measurement. Data represent mean ±SEM of n=3 replicates. Significance was determined by unpaired Student’s t-test; *P < 0.05, ****P < 0.0001. (**E**) Fluorescent micrograph of human pulmonary microvascular endothelial cells in presence and absence of IL-13 (100 ng/mL) for 48h and stained for adhesion molecules ICAM-1 and VCAM-1 (red), and VE-Cadherin (green). Nuclei were counter stained with DAPI (blue). Scale bar, 20 µm.

**Supplementary table 1.** Age and sex of donors used in this study.

| Donor # | AGE | SEX | SMOKING |
| --- | --- | --- | --- |
| 1 | 5 yo | F | N |
| 2 | 21 yo | M | N |
| 3 | 18 yo | M | Y |
| 4 | 18 yo | F | N |

**Supplementary movie 1.**
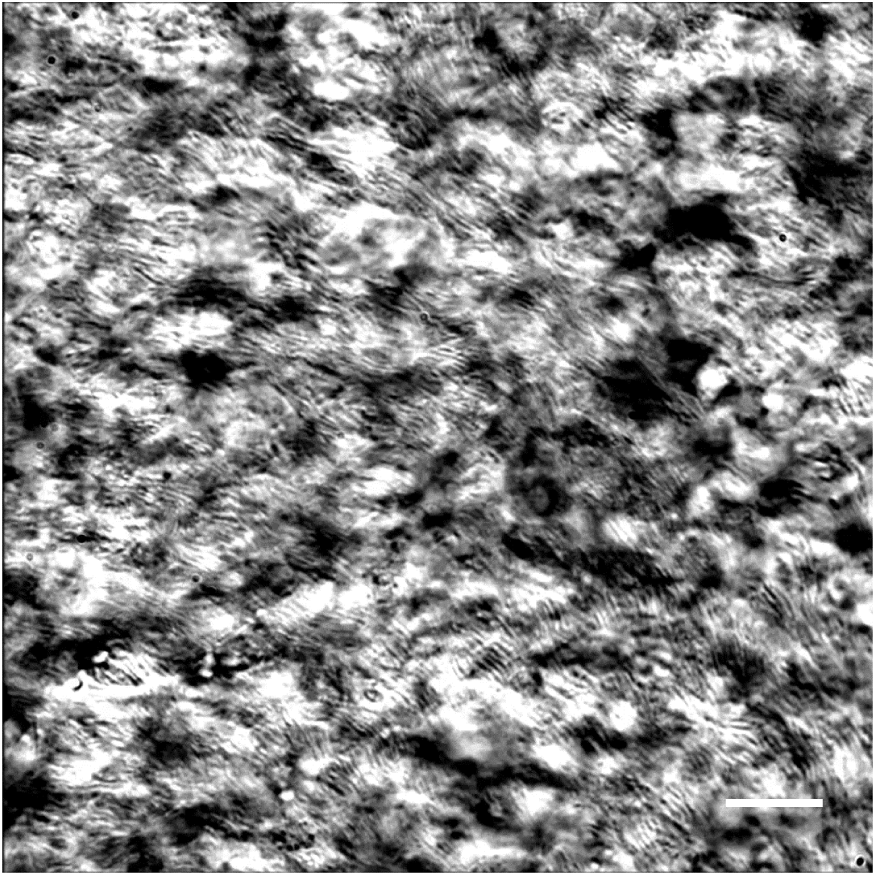
Real time phase contrast microscopic imaging of well differentiated human airway epithelial cells cultured inside the airway chip for 21 days on a 3 um pore membrane showing extensive coverage of synchronized cilia beating. Scale bar, 20 µm.

**Supplementary movie 2.**
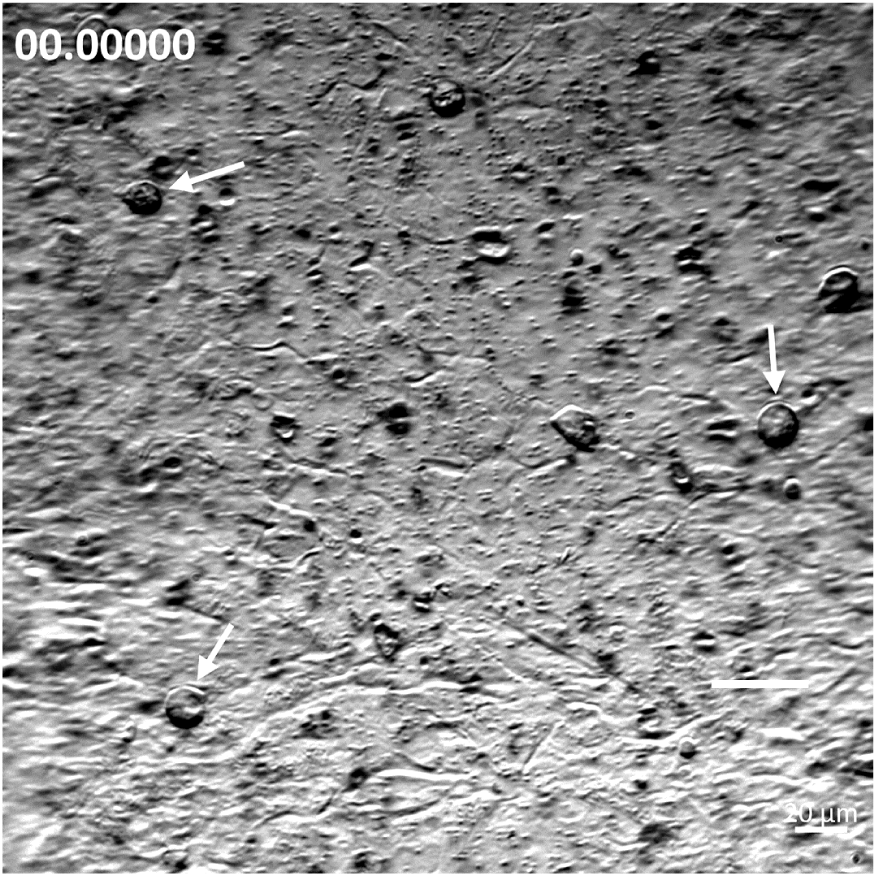
Real time phase contrast microscopic imaging of HRV16-infected airway chips (24hpi) showing rounded ciliated detaching from the epithelium (white arrows). Scale bar, 20 µm.

**Supplementary movie 3.**
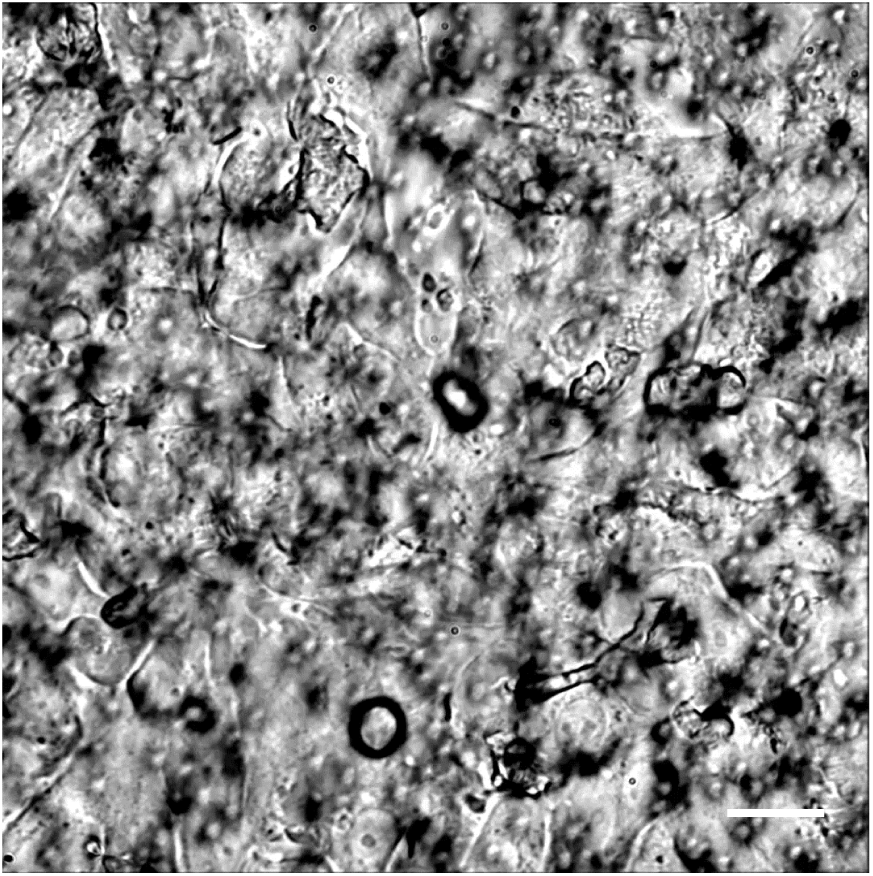
A movie showing phase contrast microscopic imaging of well differentiated human airway epithelial cells cultured inside the airway chip for 21 days on a 3 µm pore membrane and infected with HRV16 for 72 h. Scale bar, 20 µm.

**Supplementary movie 4.**
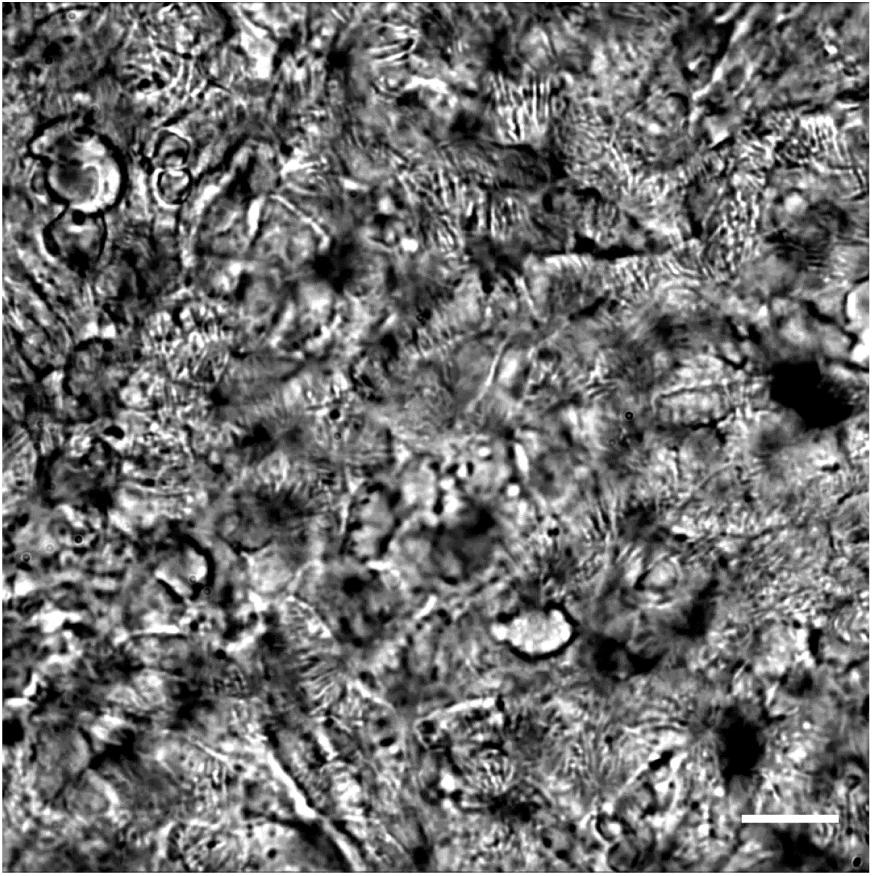
A movie showing phase contrast microscopic imaging of well differentiated human airway epithelial cells cultured inside the airway chip for 21 days on a 3 µm pore membrane and treated with IL-13 (100 ng/mL) for 7 d. Scale bar, 20 µm.

**Supplementary movie 5.**
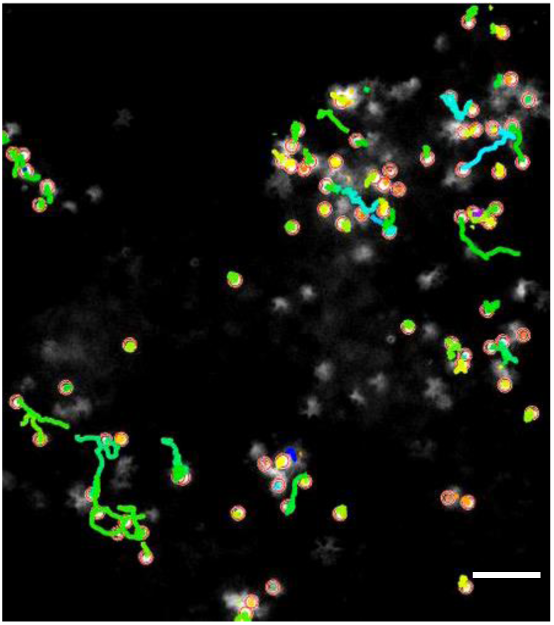
A movie showing movement tracking of fluorescently labelled human neutrophils recruited to the surface of the microvascular endothelium 24 post HRV16 infection on chip. Scale bar, 50 µm.

**Supplementary movie 6.**
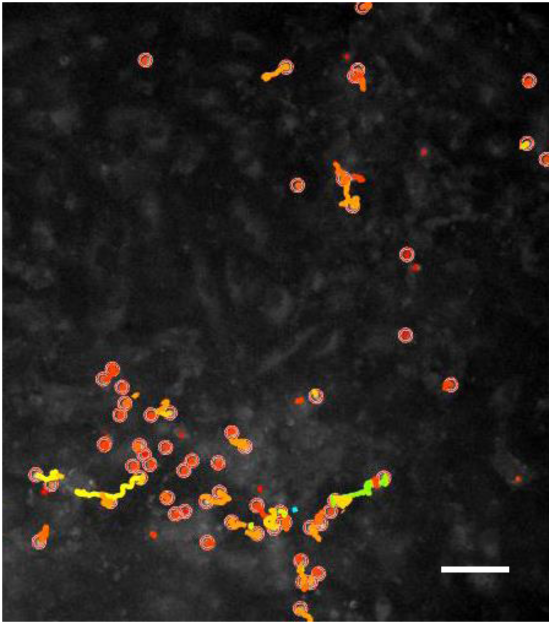
A movie showing movement tracking of fluorescently labelled human neutrophils recruited to the surface of the microvascular endothelium 24 post HRV16 infection and treated with the CXCR2 inhibitor, MK-7123 (10µM) on chip. Scale bar, 50 µm.

**Supplementary movie 7.**
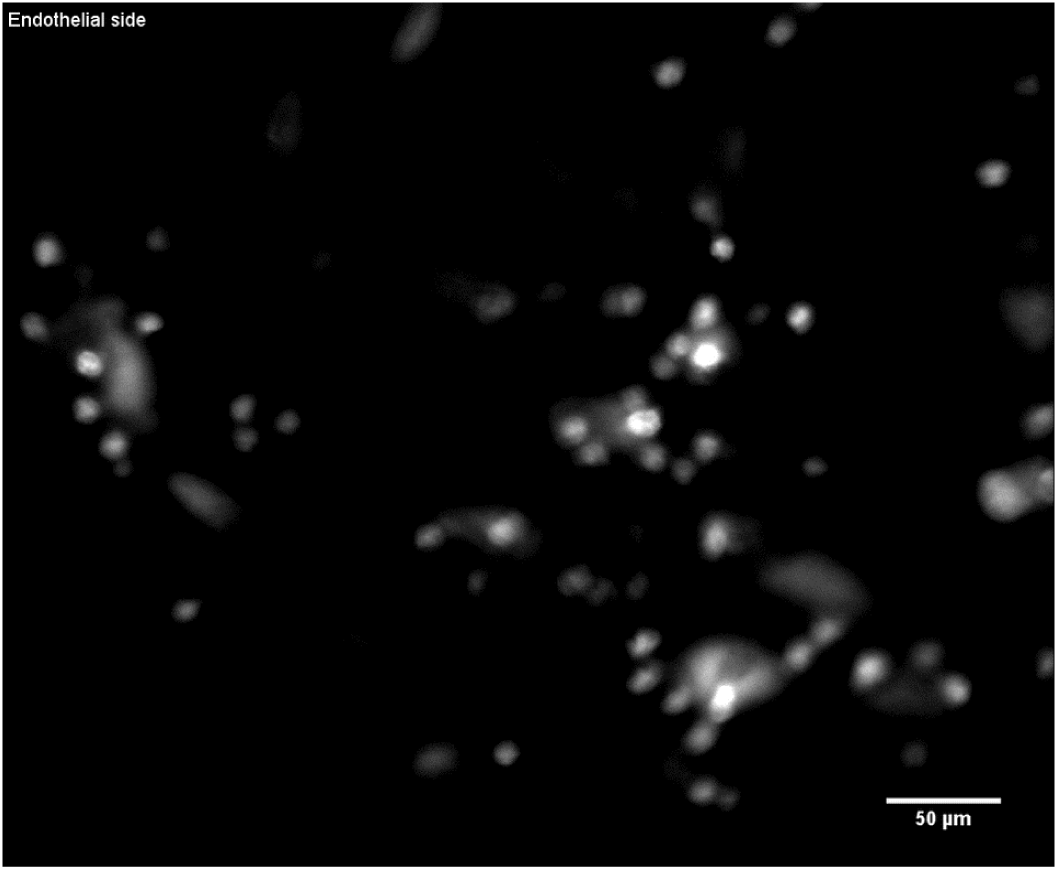
A movie of an orthogonal sectioning of a chip from the vascular to the epithelial compartment showing labelled neutrophil (white) at the surface of the microvascular endothelium and neutrophil that have crossed the membrane and transmigrated towards the epithelium. Scale bar, 50 µm.

